# EZ-FRCNN: A Fast, Accessible and Robust Deep Learning Package for Object Detection Applications from Ethology to Cell Biology

**DOI:** 10.1101/2025.06.19.660198

**Authors:** Erin Shappell, Jacob M. Wheelock, Guillaume Aubry, Hang Lu

## Abstract

Advances in high-throughput imaging and experimental automation have dramatically increased the scale of biological datasets, creating a growing need for tools that can efficiently identify and localize features in complex image data. Although deep learning has transformed image analysis, methods such as region-based convolutional neural networks remain underutilized in biology due to technical barriers such as coding requirements and reliance on cloud infrastructure. We present EZ-FRCNN, a locally hosted, user-friendly package that enables the accessible and scalable application of object detection to biological datasets. Through graphical and scriptable interfaces, users can annotate data, train models, and perform inference entirely offline. We demonstrate its utility in detecting cell phenotypes for large-scale screening, enabling the first label-free tracking of grinder motion in freely moving *C. elegans* to quantify feeding dynamics, and identifying animals in naturalistic environments for ecological field studies. These once-infeasible analyses now enable rapid screening of cell therapies, investigation of internal state–behavior coupling without immobilization or genetic modification, and efficient wildlife tracking with minimal computational cost. Together, these examples demonstrate how accessible tools like EZ-FRCNN can drive new biological discoveries in both laboratory and field environments.

## 1 Introduction

The increasing accessibility of high-resolution and high-throughput experimental methodologies has significantly expanded the amount of imaging data available to researchers in ethology and cell biology ([1–3]). In ethology, advanced imaging systems have been developed to generate vast amounts of data on model organisms such as *C. elegans* and *D. melanogaster*. These systems can capture detailed analyses of animal behavior across thousands of frames, providing unprecedented temporal resolution and enabling insights into behavioral individuality ([4]) and internal states ([5–7]). Similarly, in cellular biology, high-throughput imaging systems are used in drug screening to evaluate the efficacy of different compounds on cell viability ([8]). These systems can produce thousands of images per experiment, allowing for rapid and extensive evaluation of experimental conditions. However, generating such a large volume of images requires accessible and fast methods for data analysis, which could present challenges and therefore become the bottleneck.

While many image processing applications exist and are applicable to large datasets, accessibility remains a challenge. Many existing packages provide specialization and detailed tools that primarily benefit users with the necessary programming expertise to refine them for their problem’s needs. Two such packages are DeepLabCut (DLC) and Social LEAP Estimates Animal Poses (SLEAP), which are designed to track animal limb poses in ethological studies and offer advanced capabilities tailored to that specific application ([9–11]). However, because these packages are optimized for full-skeleton tracking, they are generally ill-suited for applications that involve tracking only a single anatomical structure. Furthermore, these programs require the use of a terminal and the setup of a virtual environment, making the setup less accessible to scientists with limited programming experience. On the other hand, CellProfiler is a comprehensive software for measuring and analyzing cell images ([12]). Its features enable detailed cell analysis but can introduce a steep learning curve for new users unfamiliar with the system. Although the complexity of these tools reflects the intricate problems they address, it can also make them less accessible to researchers who have more general needs or a less computational background.

Conversely, the rapid evolution of computer vision ecosystems can require frequent updates and changes in more user-friendly online-hosted solutions. Without constant upkeep from the original authors, these updates can lead to depreciation issues that necessitate continuous adjustments. In the worst case, unaddressed deprecation issues can break tools entirely, requiring substantial revisions to the original package. Managing broken code is particularly challenging for researchers without a computational background as it diverts valuable time and resources away from core scientific inquiries. Online hosting platforms present several other limitations. These include prohibitively expensive scaling costs for compute units and storage, code execution time limitations, and cloud compute resource unavailability, which can disrupt data processing continuity. These constraints underscore the need for robust, cost-effective, and reliable data processing solutions that can handle high-resolution and high-throughput experiments without the drawbacks of specialized software and open-source packages hosted online.

To address these challenges, we developed EZ-FRCNN, a locally run object-detection package that offers a powerful and user-friendly solution for scientists across many disciplines. This package enables researchers to quickly and reliably apply the Faster Region-based Convolutional Neural Network (FRCNN) architecture ([13]) to a wide range of images and videos, meeting the demands of increasingly high-throughput, high-resolution biological experiments. The package includes an in-house annotator for seamless data labeling and integrates a full graphical user interface (GUI) for accessibility. Rather than introducing a novel algorithm, EZ-FRCNN is designed as a resource that makes an established object detection method accessible and deployable across a range of biological contexts. We demonstrate its utility through three case studies that highlight both the versatility of the tool and the types of scientific analysis it enables. First, we demonstrate its application in cell biology by efficiently detecting and classifying adherent and detached cells in well-plate assays, where the differentiation of these phenotypes is a common requirement for high-throughput drug screening and phenotypic profiling. Next, we apply EZ-FRCNN to track the dynamics of the grinder organ in freely moving *C. elegans*, a 10 µm structure that oscillates at 4–5 Hz during feeding. This task has historically required genetic labeling, immobilization, or custom optics; here, we achieve label-free tracking in brightfield video, enabling real-time quantification of feeding dynamics and opening new avenues for studying internal state, satiety, and decision-making in unrestrained animals. Finally, we show that EZ-FRCNN can identify small dart frogs in a simulated rainforest environment, highlighting its robustness in visually complex, naturalistic conditions and its potential for field-based ecological monitoring.

## 2 Results

### 2.1 EZ-FRCNN workflow and features

EZ-FRCNN provides a streamlined workflow consisting of three main stages (Fig. 1): annotation, training, and inference. The software supports two interaction methods tailored to different user expertise levels: an intuitive graphical user interface (GUI) for users without programming experience and a customizable Jupyter notebook for users requiring more programmatic flexibility. Both methods utilize a plug-and-play approach, where data can be directly imported and processed without complicated pre-processing steps or format conversions.

**Figure 1.**
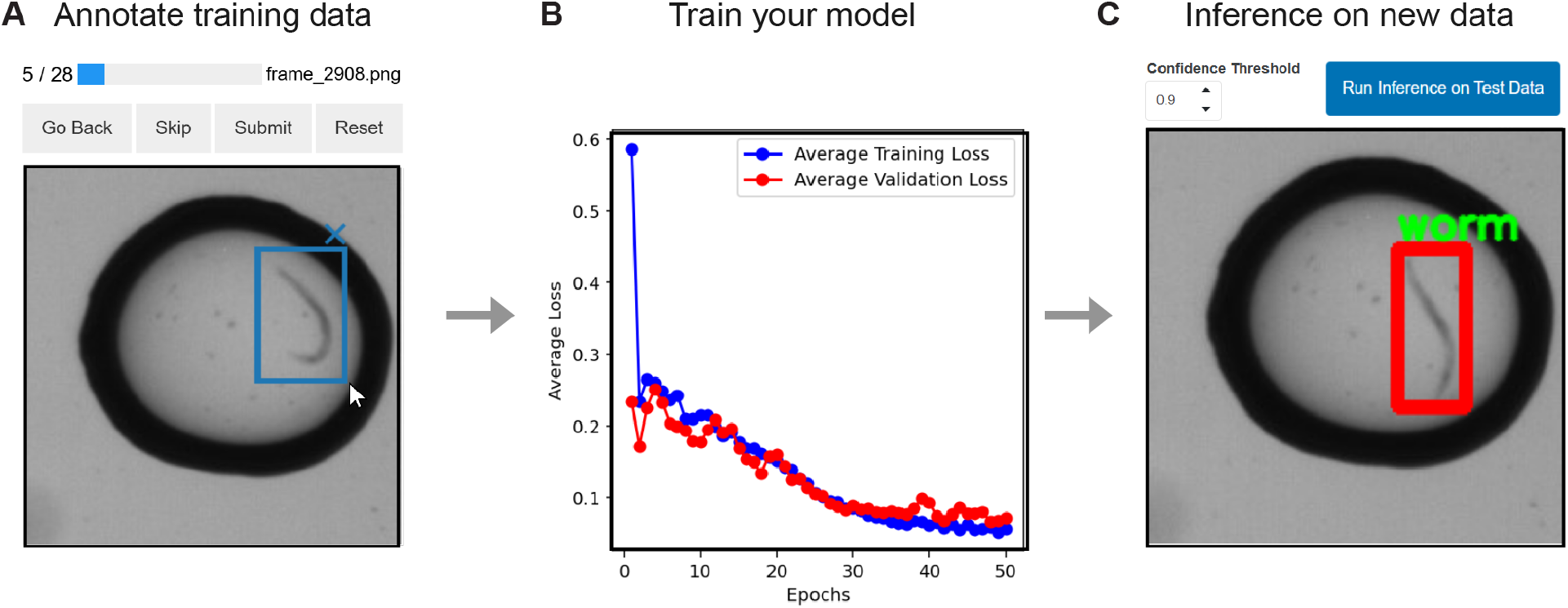
The EZ-FRCNN process is illustrated in three steps: (A) The built-in annotator allows users to label datasets using an intuitive GUI. (B) The network is trained with real-time feedback on loss metrics, making the training process easy to configure and monitor. (C) Inference is performed on new data, with visualized detection results and adjustable parameters. The entire process is designed with a user-friendly GUI to assist users through annotation, training, and inference.

In the annotation phase, users label images with minimum effort through simple mouse interactions in the GUI or through straightforward scripting in the notebook environment. During the training stage, the Faster Region-based Convolutional Neural Network (FRCNN) is trained with real-time visual feedback on loss metrics, allowing immediate adjustments to parameters like learning rate, batch size, and training duration to optimize performance. With a GPU, the final inference stage can analyze new datasets at speeds as high as 13 frames per second, providing clear visualizations of the detection results. Users can further refine analyses by adjusting detection confidence thresholds and visualization settings directly within the GUI or notebook, facilitating immediate interpretation and practical application of results.

### 2.2 Local use increases software stability and control

The selection of a local processing environment over a cloud-based platform was primarily motivated by the need to enhance the stability of the software and user control. Although cloud-based platforms such as Google Colaboratory offer accessibility, they present significant challenges that affect their reliability and longevity. Frequent environment updates on platforms such as Colab can lead to version mismatches, causing previously functional code to fail and requiring time-consuming troubleshooting, a task that may require computational expertise not readily available in all research settings. Additionally, cloud-based services often impose limitations on compute resources and session durations; for instance, Colab sessions may time out after 12 hours or less, hindering the ability to process large datasets or run long experiments without interruption.

By packaging our tool into a Docker container for local use ([14]), we effectively address these stability and control concerns. Docker encapsulates all necessary software, dependencies, and configurations, ensuring consistent performance across different computing environments without the need for extensive troubleshooting. This approach eliminates version mismatch issues caused by platform updates and grants researchers uninterrupted access to computational resources, free from session timeouts and limitations inherent to cloud services. Running the tool locally enhances reliability and efficiency, providing users with greater control over their computational environment, while remaining usable on commonly available CPU or GPU hardware.

### 2.3 Local use decreases program runtime

In addition to improving program stability, local use can significantly increase data processing speeds, addressing the critical need for efficiency in handling large datasets that are typical in scientific research. Cloud-based systems, particularly those that integrate services like Google Drive and Colaboratory, introduce substantial delays due to the overhead associated with data transfer between storage and compute resources. Studies have highlighted that data transfer bottlenecks are a significant challenge in cloud computing environments, often leading to increased latency and reduced performance efficiency ([15]). In our experience, inference times on such cloud-based systems can take tens of seconds per image, making video processing or real-time applications impractical and limiting the package’s utility.

By shifting to local processing, we have dramatically improved the processing time for EZ-FRCNN. Eliminating dependency on remote data transfers has reduced inference times to an average of 13 frames per second. This enhancement not only decreases the time from the experiment to the analysis but also increases the scope of data that can be practically processed. Furthermore, local use eliminates the need for an internet connection, making the program more accessible to field scientists who may lack access to reliable internet. With local processing, real-time or near-real-time applications become feasible, significantly expanding the package’s applicability.

### 2.4 Local use decreases recurring costs

While cloud-based platforms such as Google Colaboratory offer convenience and accessibility, they often come with limitations and subscription fees that can be prohibitive for processing large datasets or long-running experiments. For tasks that require extensive image processing or computational resources, cloud services may impose compute time restrictions and impose additional fees for access to these resources. Moreover, the use of cloud-based computing platforms often requires their storage services, further increasing costs. EZ-FRCNN addresses these concerns by executing all processing locally on personal machines. Users can leverage their existing hardware without incurring additional costs associated with cloud computing or storage. Rather than paying subscription fees and additional costs related to resource access, users can pay for the compute resources they can afford as a one-time cost. This approach provides greater control over expenses and can be particularly advantageous in educational settings, field research, or disciplines where budget constraints are more pronounced.

### 2.5 Setup and use of EZ-FRCNN is user-friendly across disciplines

Software packages intended for local installation often present challenges related to hardware variability and user expertise. Differences in operating systems, drivers, and configurations across individual machines can make setup difficult and unpredictable. These issues can be especially problematic when users lack the technical background to troubleshoot installation errors or environment conflicts. This heterogeneity poses a significant challenge, as creating multiple software versions to suit every possible setup is impractical. Our solution to this problem is the adoption of Docker. Using Docker, we package the application and all its dependencies into a container that provides a consistent environment on any hardware. This approach ensures that all users, regardless of their system specifics, can access a tool that works out of the box without requiring individual adjustments or extensive setup procedures.

On the second point, we recognize that tools like Docker and neural networks for object detection can be intimidating, especially for users unfamiliar with these technologies. To ensure that our package is accessible and usable by all, we designed it with ease of installation and use as a priority. We provide a detailed step-by-step guide that has been tested by individuals with varying levels of technical expertise, including those with minimal programming experience. Feedback from these users indicates that they were able to install and begin using the software in less than 30 minutes on average, demonstrating the approachability of our tool.

Additionally, we include a full graphical user interface (GUI) and a Jupyter Notebook within the package. The GUI allows users to interact with the software intuitively, without the need to write any code, while the Jupyter Notebook provides an interactive platform for those who prefer a code-based approach. These features help lower the barrier to entry and enable users to effectively utilize the software for their specific research needs, regardless of their prior experience with similar tools.

### 2.6 Application of EZ-FRCNN for single-cell detection

To demonstrate the utility of EZ-FRCNN in cell biology applications, we show its capability for the detection of adherent cells cultured in well plates. Well-plate assays constitute a standard approach in both academic laboratories and industrial settings (e.g. for large-scale drug discovery screens). Accurate detection of cells is typically an essential initial step in image-based assays, serving purposes such as cell counting, monitoring proliferation, and quantifying gene expression. Here, we present the application of EZ-FRCNN in analyzing bright-field microscopy images of HT1080 fibrosarcoma cells cultured in wells. The goal is to distinguish adherent cells from those that have detached—a distinction important for assessing cell viability.

We trained the neural network using 28 bright-field images, each 192×192 pixels, manually annotated to indicate 80 adherent and 80 detached cells (Fig. 2A). Post-training evaluation of network performance utilized a separate set of images containing over 100 annotations per phenotype. The network reliably detected and differentiated adherent from detached cells, as exemplified by the purple and blue markers indicating correct detections within ground-truth bounding box annotations (Fig. 2B). To quantify performance, precision, recall, and F1-score at various confidence score thresholds were calculated. The F1-score reached a maximum of 91% for adherent cells and 94% for detached cells at a threshold of 0.4 (Fig. 2C). Corresponding precision and recall values were 91% and 90% for adherent cell detection, and 92% and 96% for detached cell detection, respectively, demonstrating the network’s robust performance in accurately detecting and classifying cell phenotypes.

**Figure 2.**
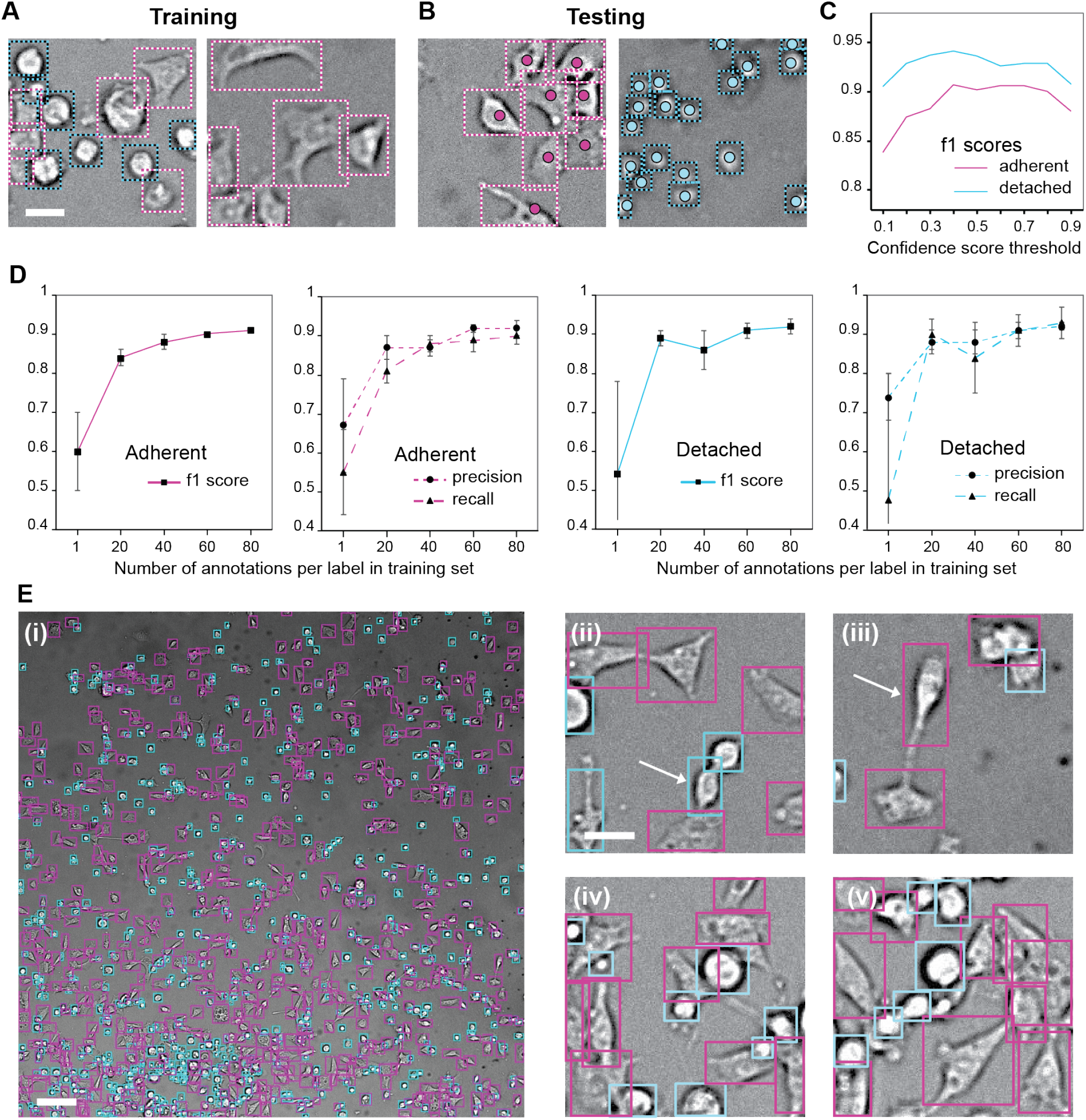
EZ-FRCNN applied to the detection of adherent cells. (a-b) Representative images used in training (a) and inferring (b). Purple/blue dashed rectangles indicate GT annotations of adherent/detached cells. Purple/blue dots indicate the center of mass of the inferences; purple for adherent cells and blue for detached cells. Scalebar is 25 µm. (c) f1 score for different confidence score thresholds used to select inferences. (d) Influence of training size on performances (f1, precision, recall). N=5 models for each data point. Error bars indicate standard deviation. (e) Representative result of a larger 2304x2304 pixel image reconstituted. (i) Whole field of view; (ii-iii) zoomed-in views. Purple/blue rectangles indicate inferences for adherent/detached cells. White arrows indicate two subtle phenotypes of adherent and detached cells. Scale bars are 100 µm in (i) and 25 µm in (ii-v).

To explore the influence of the size of the training set on overall EZ-FRCNN performance, we systematically varied the number of annotations used during training. We constructed a larger pool containing 100 annotations per phenotype and randomly sampled subsets containing 1, 20, 40, 60, or 80 annotations per class for training. For each subset size, we trained and evaluated five independent models, reporting average maximum performances (Fig. 2D). Predictably, training with only a single annotation yielded poor performance and high variability across models, as such limited data cannot encompass the diversity in image quality, cell morphology, and clustering present in the testing dataset. Notably, however, performance markedly improved with as few as 20 annotations per class, achieving F1-scores, precision, and recall above 80%, while variability decreased threefold. Performance plateaued around 90% when using 40 or more annotations, a level sufficient for many practical applications. We note that collecting this modest number of annotations is relatively rapid, requiring approximately half an hour.

To demonstrate model performance in a different context with variable cell morphologies, we analyzed a single large image containing over one thousand cells. Remarkably, the previously trained model captured the vast majority of cells, correctly categorizing their phenotypes (Fig. 2E). Note here that the pseudo ground truth is provided by a human annotator. Close examination of the annotated image reveals that there are a small number of challenging cases. First, undetected cells predominantly correspond to very low-contrast adherent cells. Their number is quantified to be less than 2%. Should higher recall be desired, the model could be further trained by supplementing the training set with low-contrast cell images. Second, many cells exhibit intermediate morphologies, where human annotators might also experience difficulty achieving consensus to binarize the morphology to be either adherent or detached (highlighted by white arrows in Fig. 2E (ii, iii)). Note that training and evaluation datasets included only cells whose adherent or detached phenotype was clearly identifiable. Interestingly, the neural network successfully detected cells of intermediate phenotype (and approximated them as either detached or adherent, or both). While human annotation forced to make a decision on either phenotypes would introduce noises, relying on automated classification presents an advantage as it would produce a consistent phenotype prediction. The cells with intermediate phenotype detected as both adherent and detached are *∼*5 % of all cells. To resolve these cases, one can simply add a criterion such as assigning the phenotype with the higher confidence score. In total, this use of EZ-FRCNN illustrates the potential for cell detection and refined phenotypic discrimination in broader applications in cell biology and drug screens.

### 2.7 EZ-FRCNN can track grinder dynamics in freely moving C. elegans

Ethological research has become increasingly dependent on high-throughput data to capture intricate details of behavior. EZ-FRCNN’s speed and accuracy in processing 4K resolution video data allow for the efficient analysis of behaviors with both large movements and fine-scale actions. To illustrate this point, we track the *C. elegans* grinder, an organ within the pharynx that breaks down food during the digestion process. The motion of the grinder is a dynamically changing behavior that can provide detailed insights into the decision making of the animal and whether the animal is actively ingesting food or not. This motion is rapid (*e*.*g*., 4–5 Hz in N2 worms) and periodic with small displacement amplitudes, necessitating a tool with high accuracy and precision for reliable tracking. Although many existing techniques have shown varying degrees of success in measuring grinder motion, these methods require fixing the worm in place, perturbing the worm with light for an extended period, or genetically modifying the worm ([16–20]). In addition, basic image processing techniques, such as thresholding, fail to track the grinder due to changes in motion and appearance in different lighting, environments, and pump stages. More advanced techniques, such as DLC and SLEAP, may underperform in tracking the grinder since these algorithms are designed to track body parts with many morphological details, which the grinder lacks. Accurately and reliably tracking the grinder requires an algorithm capable of recognizing its various appearances despite changes in lighting and position without disturbing the worm’s behavior through immobilization or genetic modification. EZ-FRCNN is a suitable algorithm for such a task. This capability enables researchers to observe both broad patterns, such as whole-body locomotion, and fine-grained, detailed behaviors, like grinder motion, simultaneously ([7, 21–25]).

Indeed, EZ-FRCNN allows for grinder tracking of freely moving worms in brightfield videos without fluorescence labeling. To meet the challenges in grinder tracking related to worm locomotion and the grinder’s size, we designed and trained a system of two EZ-FRCNN models to accomplish this task (Figure 3). First, an EZ-FRCNN model was trained to label the pharyngeal bulb (i.e., the segment of the pharynx containing the grinder). This model was trained on a set of key frames extracted from 11 videos of individual *C. elegans*. Key frames were extracted using a k-means clustering method that identifies a user-defined number (k) of the most representative frames in each video ([9, 26]). Following training and cross-validation of the pharyngeal bulb tracking model, a total of 25 videos (including the 11 used for training) were processed using the trained model. Each video was dynamically cropped to 250x250 pixels centered on the detected pharyngeal bulb to enable accurate grinder detection. The final grinder tracking EZ-FRCNN model was then trained and cross-validated by extracting a new set of key frames from the cropped versions of the same set of 11 training videos used to train the pharyngeal bulb tracking EZ-FRCNN.

**Figure 3.**
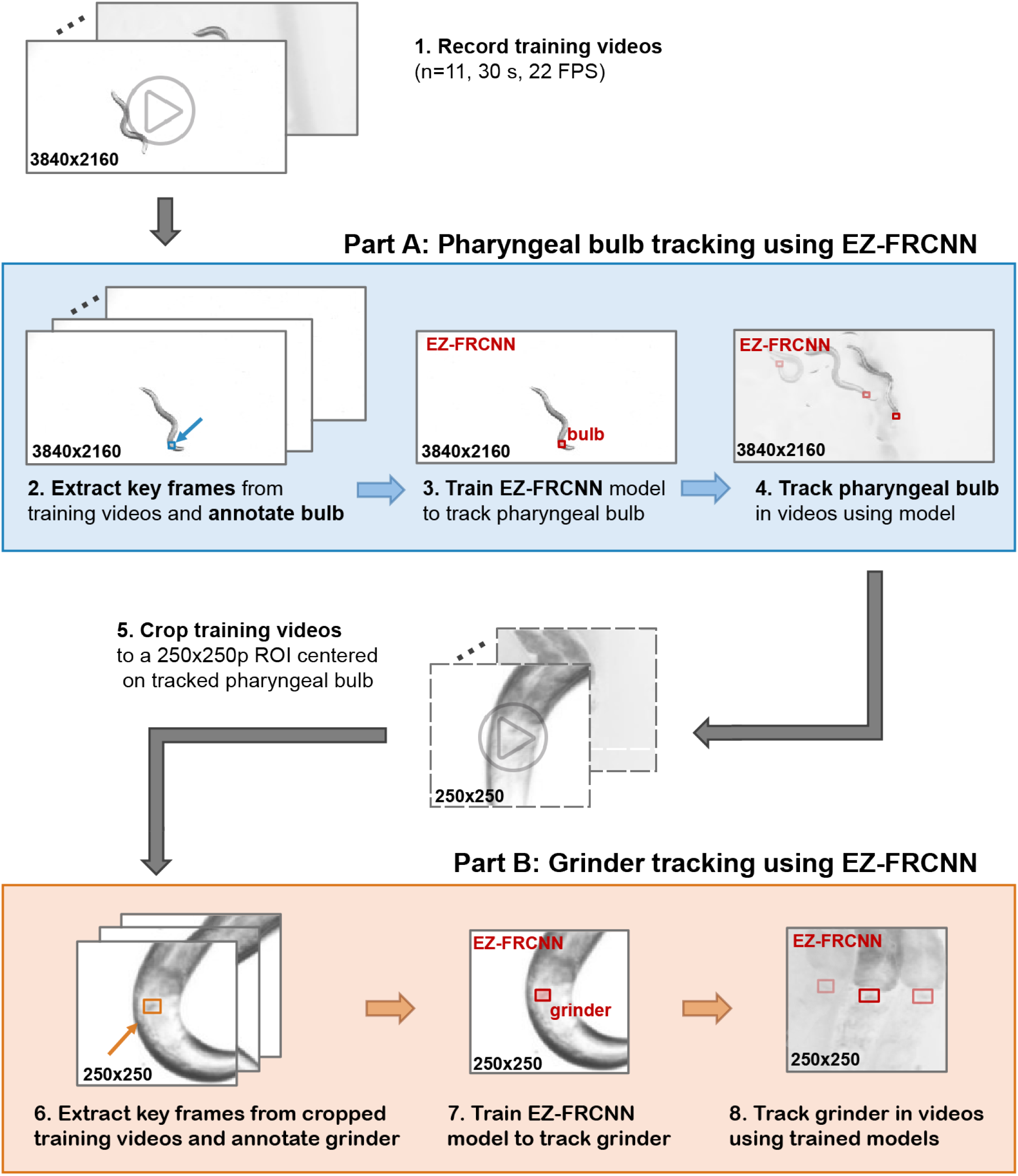
The application of two EZ-FRCNN models in series enables grinder tracking in freely moving *C. elegans*. Videos of individual worms are recorded at 22 FPS for 30 seconds each. The training data for the pharyngeal bulb tracking EZ-FRCNN is obtained by extracting 250 key frames from the videos of interest via k-means clustering. The pharyngeal bulb is annotated in all key frames and used to train the first EZ-FRCNN model. All videos are inferenced using the trained bulb tracking model, and the videos are dynamically cropped to a 250x250 pixel area centered on the center of the tracked pharyngeal bulb. After cropping the videos, 100 new key frames are extracted from the cropped videos to train the grinder tracking EZ-FRCNN. The grinder is annotated in all of the cropped key frames and used to train the second EZ-FRCNN model to track the grinder. The final model is used to track the grinder in all videos of interest.

The trained grinder tracking EZ-FRCNN is robust to blurring, contrast changes, and shape changes of the grinder (Figure 4A). This performance is made possible by training the EZ-FRCNN on examples that include roaming worms on NGM agar, dwelling worms in the bacteria lawn, and out-of-focus worms roaming on NGM agar. The number of training samples was determined via cross-validation of both the pharyngeal bulb-tracking EZ-FRCNN and the grinder-tracking EZ-FRCNN. The grinder tracking models used for cross-validation were scored on 98 key frames sampled from a separate recording across three metrics: intersection-over-union (IoU) (Figure 4B), root mean squared error (RMSE) of the tracked grinder center of mass (CoM) (Figure 4E), and dropped frame percentage (Figure 4H). In particular, there was minimal change (<1 pixel) in the RMSE of the grinder CoM as the number of training samples increased, but this is likely due to the difference in the number of grinders successfully tracked across each number of training samples. A training sample size of 100 was chosen for the final model.

**Figure 4.**
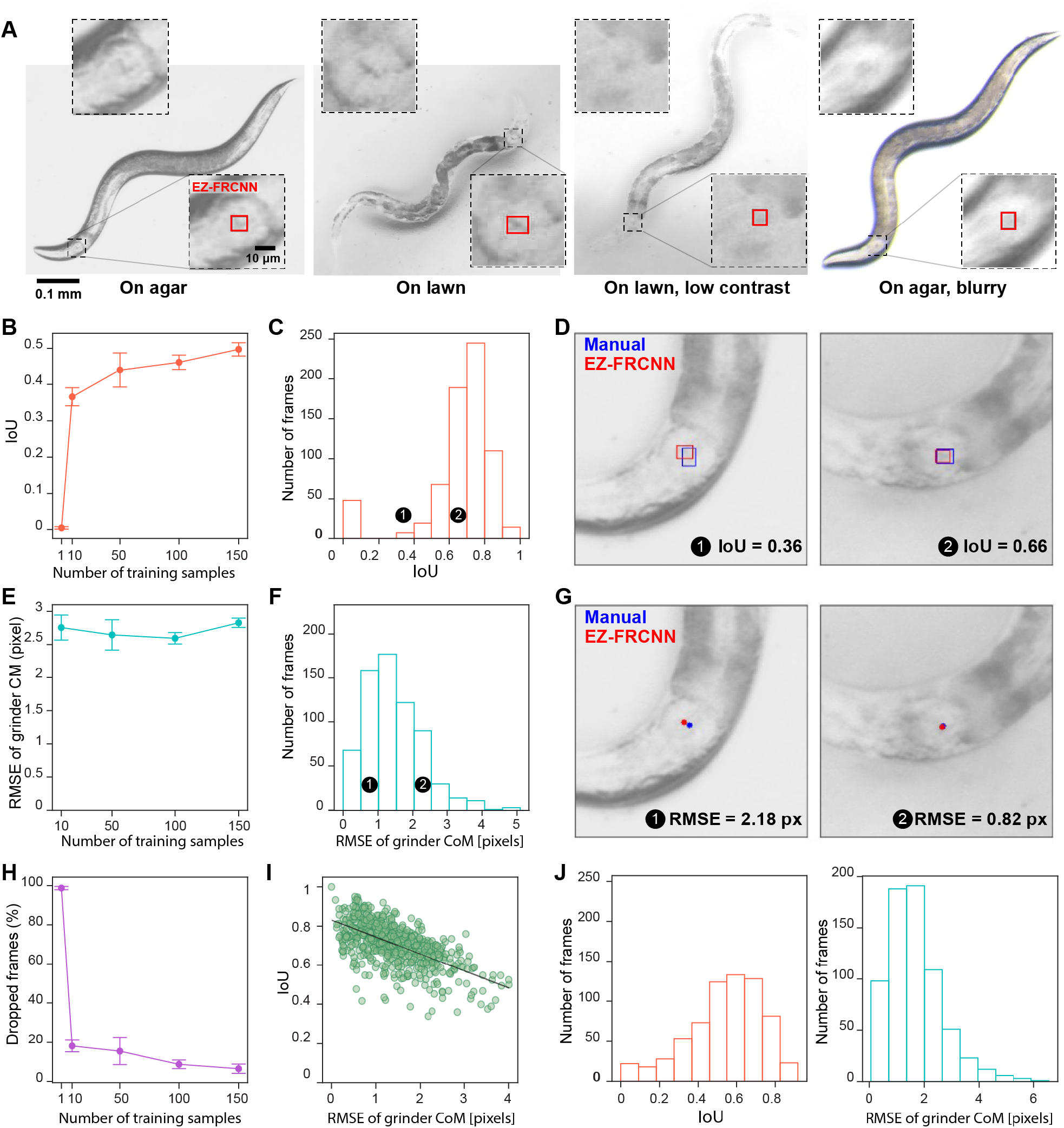
EZ-FRCNN demonstrates robust and reliable tracking of the grinder in the freely moving *C. elegans* pharynx. (A) EZ-FRCNN demonstrates robustness under various challenging conditions. The bottom inset shows a zoomed-in view with the grinder labeled by EZ-FRCNN, while the top inset shows the same view without the label. Panels BEH summarize the cross-validation performance of EZ-FRCNN on a set of 98 images when trained on bootstrapped samples of 1, 10, 50, 100, and 150 images. N=5 for error bars. (B) The intersection over union (IoU) of the EZ-FRCNN inferences compared to manual labels plateaus between 100-150 training samples. (E) There is no significant change to the root mean squared error (RMSE) of the center of mass (CoM) of the grinder for successfully tracked grinders. Note that there is no data for the 1 sample case, as no grinders were tracked successfully using this model. Average N for each number of training samples is as follows: 10 samples: 79, 50 samples: 82, 100 samples: 89, 150 samples: 91. (H) The percentage of frames dropped (i.e., not tracked) by the grinder tracking EZ-FRCNN models decreases dramatically as the sample size is increased. Panels CDFG summarize the performance of EZ-FRCNN on 700 frames sampled from 14 heldout videos after training the model on 100 frames sampled from 11 training videos. (C) Histogram of individual IoU scores between the EZ-FRCNN inferences and manual labels across all 700 frames. (D) Examples of low (left) and average (right) IoU scores. (F) Histogram of individual RMSE scores between the EZ-FRCNN-inferenced grinder CoM and those obtained using manual labels across all frames where a grinder was successfully tracked (N = 674). (G) Examples of low (left) and average (right) RMSE of grinder CoM. (I) The IoU and RMSE exhibit a strong anti-correlated relationship, indicating that a decrease in label quality (i.e., a decrease in IoU) is accompanied by an increase in grinder position tracking error (i.e., an increase in RMSE). Panel J summarizes the performance of EZ-FRCNN on tracking a full video with 683 total frames.

To quantify the performance accuracy, the final EZ-FRCNN model was used to track the grinder across a selection of 25 videos of individual *C. elegans*, 14 of which were held out from model training. Model performance on these videos was estimated by extracting 50 key frames from each video using k-means clustering, having an expert annotator manually label all key frames, and then comparing the model’s detections with the manual labels using IoU and RMSE of grinder CoM. This method for estimating video performance was used in preference to scoring every frame in every video to (1) increase the number of videos assessed and (2) to exploit the repetition of the grinder’s appearance throughout each video and reduce the number of frames necessary for scoring. However, the quantitative results for one full video (683 contiguous frames) are provided in panel J to verify that this method of using frame sampling is representative of the full video performance 4J).

Average performance across the 14 heldout videos was high, with an average of 0.66 IoU (Figure 4C), an average of less than 1.5 pixels of error in grinder position compared to manual estimates (Figure 4F), and an average of 2% dropped frames across videos. This performance exceeds that of DLC and SLEAP, whose RMSE values are generally higher than those of EZ-FRCNN (Figure 4–Supplement 1). The sample images shown in panels D and G illustrate what these metrics mean in practice: Figure 4D compares an average IoU case to an example with low IoU, which is typically considered poor, though in our data even such cases can still correspond to accurate visual tracking. Similarly, Figure 4G shows average and high RMSE examples, where higher values normally indicate poorer performance, yet often yield usable localization. Furthermore, the individual IoU and RMSE scores across all 700 key frames were shown to have a strong negative correlation, suggesting that a higher IoU score indeed suggests a more accurate grinder position track (Figure 4I). For comparison, one full video was validated, with IoU and RMSE distributions comparable to those obtained by sampling across videos 4J). Altogether, these results suggest that the tandem use of two EZ-FRCNN models can be used in series to successfully track smaller, more challenging behaviors in freely moving animals.

**Figure 4—figure supplement 1.**
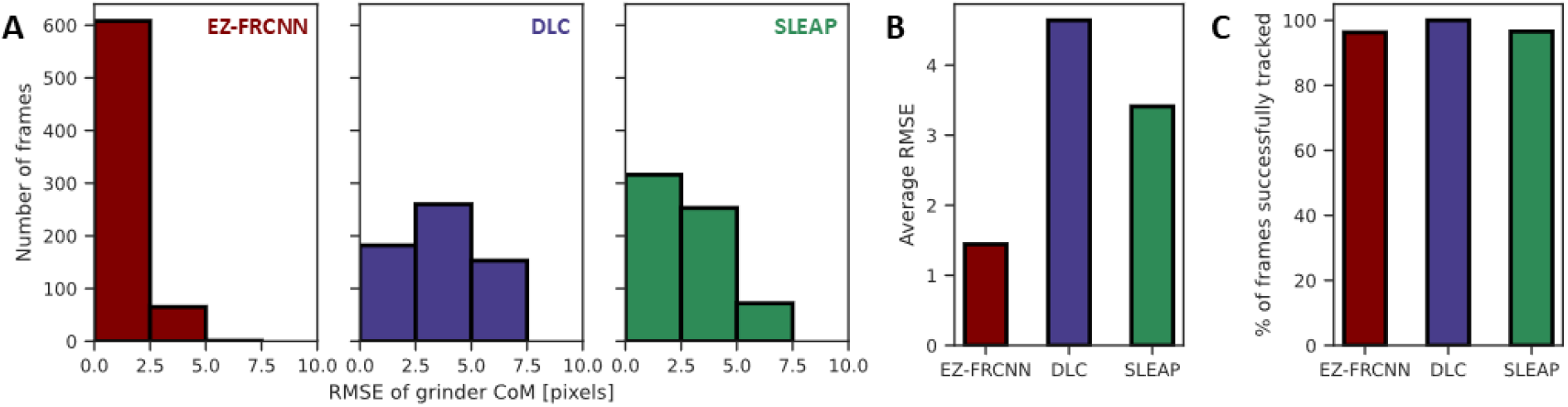
EZ-FRCNN outperforms popular pose estimation methods in single-structure tracking of the grinder in *C. elegans*. EZ-FRCNN, DLC, and SLEAP were applied to the same set of 700 frame samples used for Figure 4C,D,F,G. (A) Distribution of the RMSE of the grinder CoM for each method. RMSE is only provided for successfully tracked grinders (N=676 for EZ-FRCNN, N=700 for DLC, N=678 for SLEAP). (B) Average RMSE across methods; EZ-FRCNN is lowest. (C) Percentage of successfully tracked frames is comparable across methods.

### 2.8 Detection of Animals in Naturalistic Environments

Field ethology often requires detecting and tracking animals in environments that are visually complex, unstructured, and dynamic. These settings pose substantial challenges, including partial occlusion by vegetation, variable lighting, motion blur, and background textures that closely resemble the target organism—conditions that frequently undermine traditional image processing methods. To demonstrate how EZ-FRCNN broadens access to state-of-the-art object detection technologies in such contexts, we apply it here to the identification of Harlequin poison frogs (*O. histrionica*) in a simulated rainforest environment. Despite the heterogeneity of the scene, EZ-FRCNN reliably detects frogs even when partially obscured by foliage, as shown in Figure 5a. This result highlights the package’s ability to operate beyond controlled laboratory imaging, offering a robust solution for ecological studies in naturalistic conditions. Such adaptability is crucial for extending advanced analytical techniques to field biology, where visual noise and data limitations often impede automated analysis.

**Figure 5.**
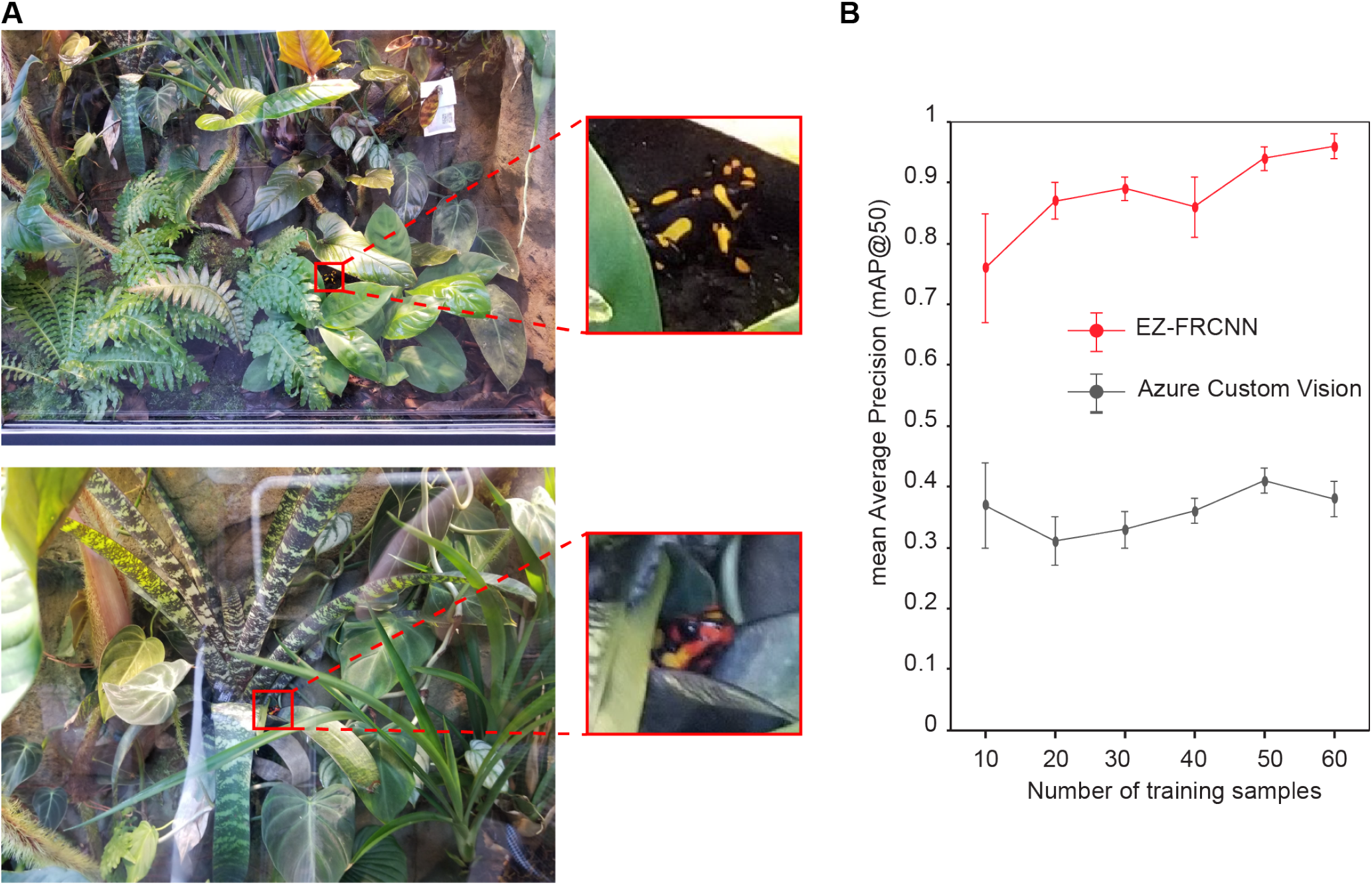
(A) Our tool’s robustness to various challenging conditions is demonstrated through successful identification of Harlequin poison frogs (*O. histrionica*) within a complex, naturalistic environment. (B) This plot illustrates the relationship between the mean validation mean average precision (mAP) and the number of training samples bootstrapped from a larger dataset. The curve shows rapid improvements with smaller sample sizes and plateaus as it approaches an optimal performance threshold, indicating the point at which additional training samples yield diminishing returns in mAP improvement.

Figure 5B further highlights EZ-FRCNN’s performance, showing a comparison in mean average precision (mAP) against Microsoft Azure Custom Vision. Mean average precision, or mAP, is a standard metric for detection quality that evaluates how accurately and consistently a model detects and locates objects. In particular, mAP@50 measures this accuracy at a 50% overlap threshold between predictions and ground truth annotations, making it a commonly used benchmark in object detection tasks. Values above 0.80 indicate strong performance in challenging conditions.

Microsoft Azure Custom Vision, a cloud-based service that enables quick, automated deployment of object detection models, is designed for users with limited technical expertise. While Azure offers ease of access, its automated approach often faces challenges in complex environments and with limited data. In contrast, EZ-FRCNN, which leverages pretraining on the COCO dataset along with extensive data augmentations, achieved a high mAP@50 of 0.90 with only 30 training images and further improved to 0.96 with 50 training images, when evaluated on a held-out set of 50 test images. Under the same conditions Azure reached only 0.41. This substantial difference demonstrates EZ-FRCNN’s capability to learn effectively from small datasets, a critical advantage for ethological studies where large-scale data gathering may be impractical.

Table 1 illustrates that EZ-FRCNN achieves commendable inference speeds on this dataset that cater well to the needs of rapid data processing in real-world scenarios. On standard GPU setups, the model processes images at an average of around 13 frames per second, facilitating near-real-time analysis, which is crucial for dynamic environmental monitoring and immediate decision-making. On CPUs, where resources are more constrained, the tool manages to maintain an average processing time of around 0.7 frames per second, ensuring broad usability even in less technologically equipped settings.

**Table 1.**
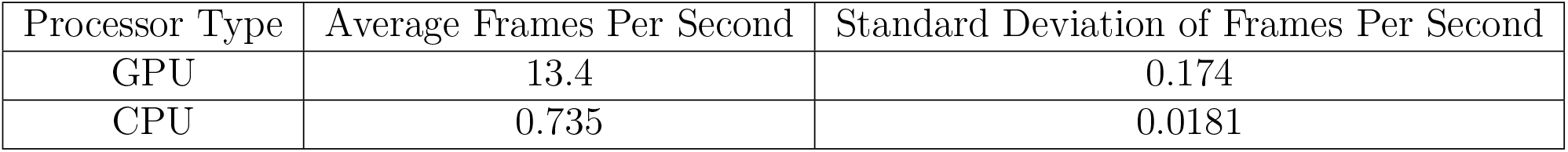
Average frames per second (FPS) and standard deviation for EZ-FRCNN inference on GPU and CPU. Higher FPS values on the GPU facilitate near-real-time image processing, while CPU performance remains adequate for less time-sensitive applications.

| Processor Type | Average Frames Per Second | Standard Deviation of Frames Per Second |
| --- | --- | --- |
| GPU | 13.4 | 0.174 |
| CPU | 0.735 | 0.0181 |

## 3 Discussion

In this study, we introduced EZ-FRCNN, a locally-run object detection package that effectively addresses the challenges of accessibility, stability, and performance inherent in existing image processing tools. By utilizing Docker for local execution, EZ-FRCNN offers a stable platform free from cloud-based dependencies. Its user-friendly graphical interface enhances accessibility for users with varying technical expertise, while leveraging local computing resources improves performance by enabling efficient processing of large datasets. Through our case studies of high-throughput cell imaging, tracking the grinder in freely moving *C. elegans*, and detecting dart frogs in natural environments, we demonstrate the tool’s robustness and scalability across diverse biological imaging applications. These examples illustrate how EZ-FRCNN successfully overcomes previous limitations, providing researchers with an effective and practical tool for advanced image analysis.

Extending this versatility to different cellular contexts, similar outcomes are anticipated when our approach to cell detection is applied to other cell lines. Many adherent cell types share common morphological features, such as distinct cellular protrusions, while suspension cells frequently resemble the detached phenotype studied here. Researchers can thus leverage our existing model readily to obtain preliminary results. Should differences in cell shape, image resolution, focus, or signal-to-noise ratio be substantial, retraining with a small dataset and optimizing confidence score thresholds based on preliminary evaluations would ensure precise and robust cell detection. We recommend following these practical guidelines to facilitate the broad and effective utilization of EZ-FRCNN in diverse well-plate assays across different cell types.

Through our application in grinder tracking in *C. elegans*, we demonstrate that EZ-FRCNN enables new forms of biological measurement—specifically, the real-time, label-free quantification of feeding behavior in unrestrained animals. To accomplish this, we used two EZ-FRCNN models in tandem: one to localize the pharyngeal region and another to detect the grinder, a 10 µm structure with low visual contrast that is easily confounded with nearby anatomy. This two-step detection strategy is broadly applicable to other systems where fine-scale motor features—such as pupil dilation or whisker movement—must be isolated within a dynamic background. By supporting this workflow without specialized software or extensive customization, EZ-FRCNN provides a flexible, accessible solution for tracking subtle behaviors across diverse biological contexts.

This example also illustrates a broader point about the utility of object detection. While pixel-wise segmentation methods like U-Net are often favored for their spatial precision, they are not always practical in situations where target structures have variable appearance, weak contrast, or ambiguous boundaries. In contrast, bounding box–based object detectors, such as Faster R-CNN, can offer more robust performance in these cases by focusing on the reliable localization of features rather than pixel-level delineation. In our grinder-tracking application, segmentation-based approaches fail to generalize across lighting and posture changes, whereas detection reliably captures the grinder’s location frame-to-frame. Detection can also serve as a precursor to segmentation by narrowing the region of interest, enabling hybrid pipelines. EZ-FRCNN makes this flexible detection-based strategy readily accessible, particularly in use cases where segmentation pipelines are brittle or infeasible.

Beyond these immediate applications, EZ-FRCNN also has potential for deployment in field-based ecological research. With additional optimization through model pruning and quantization, the tool could be adapted to run on lightweight embedded systems, enabling real-time object detection in remote or resource-limited environments without the need for high-power computing or internet access. This would allow continuous, on-site monitoring of animal populations using compact and autonomous systems. For instance, in studies of amphibian behavior, EZ-FRCNN could support long-term, real-time tracking of individual frogs in natural habitats—tasks that are difficult with cloud-based pipelines or manual video review. Access to such data is critical for understanding habitat use, movement patterns, and environmental responses, and could directly inform conservation strategies by identifying critical microhabitats or migration corridors ([27, 28]). By bringing advanced detection capabilities into the field, EZ-FRCNN expands the range of ecological questions that can be addressed with modern imaging approaches.

Another promising extension of EZ-FRCNN lies in precision agriculture, where real-time monitoring of crop health, pest presence, and growth dynamics is essential for effective farm management. By running on locally captured images from handheld devices or field cameras, EZ-FRCNN can enable continuous, in-field analysis without requiring cloud connectivity or specialized computing. This accessibility makes advanced image analysis feasible for farmers and agricultural researchers without machine learning expertise. For example, the tool could detect early signs of nutrient deficiency, disease, or pest infestation—conditions that often require prompt intervention to prevent yield loss. Rapid, on-site identification of such stressors allows for more targeted and efficient resource use. By facilitating timely, data-driven decisions, EZ-FRCNN supports sustainable agricultural practices and improves crop productivity across a range of farming contexts ([29]).

Overall, EZ-FRCNN addresses a critical gap between the increasing volume of biological imaging data and the accessibility of advanced analytical tools. By lowering technical barriers, the package empowers a broader range of researchers to perform object detection tasks across diverse experimental contexts, from lab-based assays to field deployments. The case studies and applications we present demonstrate that this accessibility is not merely a convenience, but a scientific enabler that allows researchers to quantify behaviors, phenotypes, and environmental interactions that were previously difficult or impossible to measure. As imaging technologies continue to proliferate across biology, tools like EZ-FRCNN will play a central role in unlocking the full scientific potential of these datasets. By continuing to refine and extend this platform, we aim to support a more inclusive and insightful era of image-based biological discovery.

## 4 Methods and Materials

### 4.1 C. elegans maintenance

All *C. elegans* strains were cultured on Nematode Growth Media (NGM) plates at 20°C. NGM plates were seeded with 200 µL of OP50 (cultured overnight, in LB). The strains used in this work include N2, DA1113 [*eat-2 (ad1113)*], and MT10549 [*tdc-1(n3421)*].

### 4.2 C. elegans recordings

All *C. elegans* recordings were completed using a stereomicroscope (Leica M165 FC) with an 8.3 megapixel camera (InfinityCam 8-8). Worms were transferred via pipette to NGM plates either seeded with OP50 or NGM plates without OP50. After 15 minutes, 5 random worms were selected from each plate and recorded in 4K resolution at 6.3x magnification for 30 seconds at an acquisition rate of 22 FPS.

### 4.3 Cell maintenance

Cell assays were performed using adherent epithelial HT-1080 cell line (ATCC). The HT-1080 cells were cultured in tissue culture flasks using DMEM medium (ATCC) supplemented with 10% fetal bovine serum albumin and 100 units/mL penicillin streptomycin (VWR). Cells were cultured at 37 °C in a humidified 5 % CO2 incubator.

### 4.4 Cell detection assay

For cell detection assays, cells were trypsinized with 0.25% trypsin-EDTA and resuspended in cell culture medium at 0.01x10^6^ cells /mL. 100 uL were seeded in the wells of a 96 well plate. Microscopy images were taken at 0 hour, 4 hours, and 15 hours after seeding using a Nikon W1 spinning disk confocal system.

### Hardware setup for timing analysis

All timing experiments in this study were conducted on a system configured with the following specifications:

- **Processor**: AMD Ryzen 9 5900X 12-Core Processor, running at 3.70 GHz
- **Memory**: 64 GB RAM
- **GPU**: NVIDIA GeForce RTX 4070 Ti

### EZ-FRCNN characterization

The EZ-FRCNN model is based on the Faster R-CNN architecture ([13]), which includes the following key components:

1. **Backbone (ResNet-50 with FPN)**: A pre-trained ResNet-50 model with a Feature Pyramid Network (FPN) is used to extract multi-scale feature maps from input images. This enables effective detection of objects at various scales.
2. **Region Proposal Network (RPN)**: The RPN slides over the feature maps generated by the backbone to propose regions of interest (ROIs) that likely contain objects. Each region is associated with a confidence score.
3. **ROI Pooling and Head Network**: Proposed ROIs are processed through an ROI pooling layer to standardize the feature map size, then fed into fully connected layers that predict class labels and refine bounding box coordinates.
4. **Loss Function**: The total loss *L* for Faster R-CNN combines classification and regression losses to optimize both object classification and localization. It is defined as:

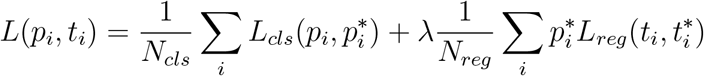

where:

- *i* indexes the anchor or proposal in the mini-batch.
- *p*_*i*_ is the predicted probability of the anchor *i* being an object.
- *p*^*∗*^_*i*_ is the ground truth label (1 if the anchor is positive, 0 if negative).
- *t*_*i*_ is the predicted bounding box coordinates for anchor *i*.
- *t*^*∗*^_*i*_ is the ground truth bounding box coordinates for anchor *i*.
- *N*_*cls*_ is the number of anchors (or proposals) in the mini-batch for the classification loss.
- *N*_*reg*_ is the number of anchors (or proposals) in the mini-batch for the regression loss (usually set to 1 for convenience).
- *λ* is a balancing parameter to weigh the two loss components, typically set to *λ* = 1.

The classification loss *L*_*cls*_ is a log loss over two classes (object vs. background) and is defined as:

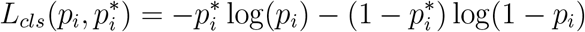

The regression loss *L*_*reg*_ is a smooth L1 loss function between the predicted bounding box coordinates and the ground truth bounding box coordinates, given by:

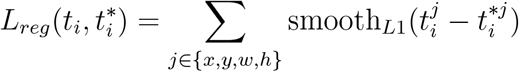

where the smooth L1 loss is defined as:

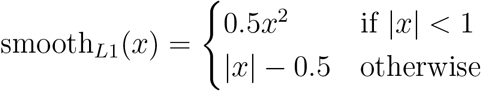

### EZ-FRCNN training

The training process uses a standard pipeline for optimizing model weights, detailed as follows:

1. **Initialization**: The Faster R-CNN model with a ResNet-50 backbone is loaded with COCO-pretrained weights. The classification layer (head) is modified to match the specific number of classes for the task.
2. **Optimizer**: Stochastic Gradient Descent (SGD) is used with a learning rate of 0.001, momentum of 0.9, and weight decay of 0.0005.
3. **Training Loop**: For each epoch, the model iterates over the training data to compute and backpropagate losses, updating weights to minimize a combined classification and bounding box regression loss.
4. **Validation**: After each epoch, the model is evaluated on a validation dataset to monitor performance and prevent overfitting.
5. **Checkpointing**: Model weights and loss curves are saved at regular intervals to facilitate training monitoring and recovery.

### EZ-FRCNN inferencing

The inferencing process enables detection of objects in new images or videos, leveraging the trained model’s predictive capabilities:

**1. Model Loading and Evaluation Mode**: The trained weights are loaded, and the model is set to evaluation mode to ensure deterministic predictions.

**2. Detection on Images or Videos**: The model takes in images or video frames, generating bounding boxes, class labels, and confidence scores for each detected object. Only detections above a user-defined confidence threshold are retained.

**3. Drawing Detections and Saving Results**: For each detected object, bounding boxes and labels are drawn onto the image or video frame, and the results are saved in the output directory. Bounding box coordinates, class labels, and confidence scores are also logged in a CSV file.

### Cropping videos using tracked pharyngeal bulb location

One of the challenges of tracking a small moving structure within a free moving organism is obtaining a region of interest (ROI) that eliminates false positives that may be found throughout the organism. In our application to *C. elegans*, we exploited the consistent appearance of the pharyngeal bulb, the part of the pharynx that contains the structure of interest, in this task. We trained an EZ-FRCNN model to track the pharyngeal bulb, then used the center of the resulting bounding box from each frame to dynamically crop each video to a 250 x 250 pixel ROI centered on the pharyngeal bulb. Before frame cropping, the bounding box centers are interpolated and smoothed using a Gaussian filter to fill in dropped frames and ensure a smooth ROI video.

### Optimizing key frame selection with k-means clustering

Training data diversity is an important factor in training a generalizable machine learning algorithm, including one such as EZ-FRCNN. To streamline the process of selecting the most diverse set of frames from training videos, we used DeepLabCut’s implementation of k-means clustering ([9, 26]). In contrast to a uniform sampling method, a k-means sampling method will take all frames from a video, sort them into (k) clusters based on similarity, and then select one frame from each cluster to use for training. This ensures that the final training set includes all possible appearances of the structure being tracked and improves model performance.

### Optimizing training set size through cross-validation

Another essential feature in training is the size of the training set. A training set that is too small may produce a model incapable of performing on heldout data, while a training set that is too large is more prone to overfitting. To find the ideal training set size for the two EZ-FRCNN models trained to track the *C. elegans* pharyngeal bulb and the grinder, cross-validation was completed for five different training set sizes.

Sample sizes of 1, 10, 50, 100, and 250 were used for the pharyngeal bulb tracking model, while sizes of 1, 10, 50, 100, and 150 were used for the grinder tracking model. For each sample size, five random shuffles were bootstrapped from the same set of 499 samples for the pharyngeal bulb tracking model and 192 for the grinder tracking model. Training sets were sampled via k-means from a recording containing 19 videos of individual *C. elegans*. Each bootstrapped model was then tested on a heldout set of 98 frames sampled via k-means from a separate recording of 12 videos. The frame size used for pharyngeal bulb cross-validation was the original frame size of 3840 x 2160 pixels, while the grinder cross-validation images were 250 x 250 pixel cropped sections of these frames that were centered on the pharyngeal bulb. The bootstrapped models of each sample size were scored using intersection over union (IoU), root mean squared error (RMSE) of the grinder center of mass (CoM), and the percentage of frames dropped (i.e., not tracked). 4bcd summarize the results for each sample size.

## Data availability

Data supporting the findings of this study are available upon request or at https://osf.io/z7t9s.

## Code availability

EZ-FRCNN is available at https://github.com/lu-lab/ez-frcnn and www.ezfrcnn.com. Docker images and usage instructions are included.

## Acknowledgements

This material is based upon work supported by the National Science Foundation Graduate Research Fellowship (NSF GRFP) under Grant No. DGE-2039655 to ES, the Integrative and Quantitative Biosciences Accelerated Training Environment (InQuBATE) T32 program funded by the National Institute of General Medical Sciences (NIGMS) under Grant No. T32GM142616 to JMW, the National Science Foundation (NSF) under Grant No. 1648035 to GA, the National Institute of Medical Sciences (NIGMS) under Grant No. T32GM142616, NSF 1764406, and NIH R01AG082039 to HL.

## Author contributions

E.S. and J.M.W. contributed equally to development, experiments, and manuscript writing. G.A. supervised biological validations. H.L. supervised the project and provided critical feedback.

## References

[1] Kathrin Steck, Daniel Veit, Ronald Grandy, Sergi Bermúdez i Badia, Zenon Mathews, Paul F. M. J. Verschure, Bill S. Hansson, and Markus Knaden. A high-throughput behavioral paradigm for drosophila olfaction -the flywalk. Scientific Reports, 2012. doi: 10.1038/srep00361.

[2] Matthew A. Churgin, Sang Kyu Jung, Chih Chieh Yu, Xiangmei Chen, David M. Raizen, and Christopher Fang-Yen. Longitudinal imaging of caenorhabditis elegans in a microfabricated device reveals variation in behavioral decline during aging. eLife, 2017. doi: 10.7554/elife.26652.

[3] Kim Hung Le, Min Zhan, Yong-Min Cho, Jason Wan, Jason Wan, Dhaval Patel, and Hang Lu. An automated platform to monitor long-term behavior and healthspan in caenorhabditis elegans under precise environmental control. Communications Biology, 2020. doi: 10.1038/s42003-020-1013-2.

[4] Kyle S. Honegger, Matthew A.-Y. Smith, Matthew A. Churgin, Glenn C. Turner, and Benjamin L. de Bivort. Idiosyncratic neural coding and neuromodulation of olfactory individuality in ¡i¿drosophila¡/i¿. Proceedings of the National Academy of Sciences, 117 (38):23292–23297, 2020. doi: 10.1073/pnas.1901623116. URL https://www.pnas.org/doi/abs/10.1073/pnas.1901623116.

[5] David J. Anderson. Circuit modules linking internal states and social behaviour in flies and mice. Nature Reviews Neuroscience, 17(11):692–704, October 2016. ISSN 1471-0048. doi: 10.1038/nrn.2016.125. URL http://dx.doi.org/10.1038/nrn.2016.125.

[6] Steven W. Flavell, Nadine Gogolla, Matthew Lovett-Barron, and Moriel Zelikowsky. The emergence and influence of internal states. Neuron, 110(16):2545–2570, August 2022. ISSN 0896-6273. doi: 10.1016/j.neuron.2022.04.030. URL http://dx.doi.org/10.1016/j.neuron.2022.04.030.

[7] Elias Scheer and Cornelia I Bargmann. Sensory neurons couple arousal and foraging decisions in Caenorhabditis elegans. eLife, 12:RP88657, Dec 2023. ISSN 2050-084X. doi: 10.7554/eLife.88657. URL https://doi.org/10.7554/eLife.88657.

[8] Mark-Anthony Bray, Shantanu Singh, Han Han, Chadwick T. Davis, Sadhna Phanse, Blake Borgeson, Cathy L Hartland, Maria Kost-Alimova, Sigrún M. Gústafsdóttir, Sigrun M Gustafsdottir, Christopher C. Gibson, and Anne E. Carpenter. Cell painting, a high-content image-based assay for morphological profiling using multiplexed fluorescent dyes. Nature Protocols, 2016. doi: 10.1038/nprot.2016.105.

[9] Tanmay Nath, Alexander Mathis, An Chi Chen, Amir Patel, Matthias Bethge, and Mackenzie Weygandt Mathis. Using deeplabcut for 3d markerless pose estimation across species and behaviors. Nature Protocols, 14(7):2152–2176, June 2019. ISSN 1750-2799. doi: 10.1038/s41596-019-0176-0. URL http://dx.doi.org/10.1038/s41596-019-0176-0.

[10] Jessy Lauer, Mu Zhou,, Shaokai Ye,, William Menegas, Steffen Schneider, Tanmay Nath, Mohammed Mostafizur Rahman, Valentina Di Santo, Daniel Soberanes, Guoping Feng, Venkatesh N. Murthy, George Lauder, Catherine Dulac, Mackenzie Weygandt Mathis, Mackenzie W. Amoroso, and Alexander Mathis. Multi-animal pose estimation, identification and tracking with deeplabcut. Nature Methods, 2022. doi: 10.1038/s41592-022-01443-0.

[11] Talmo D. Pereira, Nathaniel Tabris, Arie Matsliah, David M. Turner, Junyu Li, Shruthi Ravindranath, Eleni S. Papadoyannis, Edna Normand, David S. Deutsch, Z. Yan Wang, and et al. Sleap: A deep learning system for multi-animal pose tracking. Nature Methods, 19(4):486–495, Apr 2022. doi: 10.1038/s41592-022-01426-1.

[12] David R. Stirling, Madison J. Swain-Bowden, Alice M. Lucas, Anne E. Carpenter, Beth A. Cimini, and Allen Goodman. Cellprofiler 4: improvements in speed, utility and usability. BMC Bioinformatics, 2021. doi: 10.1186/s12859-021-04344-9.

[13] Shaoqing Ren, Kaiming He, Ross Girshick, and Jian Sun. Faster r-cnn: Towards real-time object detection with region proposal networks. In C. Cortes, N. Lawrence, D. Lee, M. Sugiyama, and R. Garnett, editors, Advances in Neural Information Processing Systems, volume 28. Curran Associates, Inc., 2015. URL https://proceedings.neurips.cc/paper_files/paper/2015/file/14bfa6bb14875e45bba028a21ed38046-Paper.pdf.

[14] Dirk Merkel. Docker: lightweight linux containers for consistent development and deployment. Linux journal, 2014(239):2, 2014.

[15] Chaowei Yang, Qunying Huang, Zhenlong Li, Kai Liu, and Fei Hu. Big data and cloud computing: Innovation opportunities and challenges. International Journal of Digital Earth, 10(1):13–53, Nov 2016. doi: 10.1080/17538947.2016.1239771.

[16] David Raizen. Methods for measuring pharyngeal behaviors. WormBook, page 1–13, December 2012. ISSN 1551-8507. doi: 10.1895/wormbook.1.154.1. URL http://dx.doi.org/10.1895/wormbook.1.154.1.

[17] Monika Scholz, Dylan J. Lynch, Kyung Suk Lee, Erel Levine, and David Biron. A scalable method for automatically measuring pharyngeal pumping in c. elegans. Journal of Neuroscience Methods, 274:172–178, 2016. ISSN 0165-0270. doi: 10.1016/j.jneumeth.2016.07.016. URL https://www.sciencedirect.com/science/article/pii/S0165027016301777.

[18] Kyung Suk Lee, Shachar Iwanir, Ronen B. Kopito, Monika Scholz, John A. Calarco, David Biron, and Erel Levine. Serotonin-dependent kinetics of feeding bursts underlie a graded response to food availability in c. elegans. Nature Communications, 8(1), February 2017. ISSN 2041-1723. doi: 10.1038/ncomms14221. URL http://dx.doi.org/10.1038/ncomms14221.

[19] Ma Jesús Rodríguez-Palero, Ana López-Díaz, Roxane Marsac, José-Eduardo Gomes, María Olmedo, and Marta Artal-Sanz. An automated method for the analysis of food intake behaviour in caenorhabditis elegans. Scientific Reports, 8(1), February 2018. ISSN 2045-2322. doi: 10.1038/s41598-018-21964-z. URL http://dx.doi.org/10.1038/s41598-018-21964-z.

[20] Elsa Bonnard, Jun Liu, Nicolina Zjacic, Luis Alvarez, and Monika Scholz. Automatically tracking feeding behavior in populations of foraging C. elegans. eLife, 11:e77252, sep 2022. ISSN 2050-084X. doi: 10.7554/eLife.77252. URL https://doi.org/10.7554/eLife.77252.

[21] Juliette Ben Arous, Sophie Laffont, and Didier Chatenay. Molecular and sensory basis of a food related two-state behavior in c. elegans. PLOS ONE, 4(10):1–8, 10 2009. doi: 10.1371/journal.pone.0007584. URL https://doi.org/10.1371/journal.pone.0007584.

[22] Karl Emanuel Busch and Birgitta Olofsson. Should i stay or should i go? Worm, 1 (3):182–186, 2012. doi: 10.4161/worm.20464. URL https://doi.org/10.4161/worm.20464. PMID: 24058845.

[23] Monika Scholz, Aaron R. Dinner, Erel Levine, and David Biron. Stochastic feeding dynamics arise from the need for information and energy. Proceedings of the National Academy of Sciences, 114(35):9261–9266, 2017. doi: 10.1073/pnas.1703958114. URL https://www.pnas.org/doi/abs/10.1073/pnas.1703958114.

[24] Steven W Flavell, David M Raizen, and Young-Jai You. Behavioral States. Genetics, 216(2):315–332, 10 2020. ISSN 1943-2631. doi: 10.1534/genetics.120.303539. URL https://doi.org/10.1534/genetics.120.303539.

[25] Adam A Atanas, Jungsoo Kim, Ziyu Wang, Eric Bueno, Mccoy Becker, D. Kang, Jungyeon Park, Talya S Kramer, Flossie K Wan, Saba Baskoylu, Ugur Dag, Elpiniki Kalogeropoulou, Matthew A Gomes, Cassi Estrem, Netta Cohen, Vikash K Mansinghka, and Steven W Flavell. Brain-wide representations of behavior spanning multiple timescales and states in c. elegans. Cell, 186(19):4134–4151.e31, September 2023.

[26] Alexander Mathis, Pranav Mamidanna, Taiga Abe, Kevin M. Cury, Venkatesh N. Murthy, Mackenzie Weygandt Mathis, and Matthias Bethge. Markerless tracking of user-defined features with deep learning. arXiv: Computer Vision and Pattern Recognition, 2018. doi: 10.1038/s41593-018-0209-y.

[27] Erin Muths, Alisa L. Gallant, Evan H. Campbell Grant, William A. Battaglin, David E. Green, Jennifer S. Staiger, Susan C. Walls, Margaret S. Gunzburger, and Rick F. Kearney. The amphibian research and monitoring initiative (armi): 5-year report. Scientific Investigations Report, 2006. doi: 10.3133/sir20065224.

[28] Andrew J. Hamer, Andrew J. Hamer, Mark J. McDonnell, and Mark J. McDonnell. Amphibian ecology and conservation in the urbanising world: A review. Biological Conservation, 2008. doi: 10.1016/j.biocon.2008.07.020.

[29] Konstantinos G Liakos, Patrizia Busato, Dimitrios Moshou, Simon Pearson, and Dionysis Bochtis. Machine learning in agriculture: A review. Sensors, 2018. doi: 10.3390/s18082674.

